# Instability of excitatory synapses in experimental autoimmune encephalomyelitis and the outcome for excitatory circuit inputs to individual cortical neurons

**DOI:** 10.1101/2024.01.23.576662

**Authors:** Rebecca L. Gillani, Eseza N. Kironde, Sara Whiteman, Theodore J. Zwang, Brian J. Bacskai

## Abstract

Synapses are lost on a massive scale in the brain and spinal cord of people living with multiple sclerosis (PwMS), and this synaptic loss extends far beyond demyelinating lesions. Post-mortem studies show the long-term consequences of multiple sclerosis (MS) on synapses but do not inform on the early impacts of neuroinflammation on synapses that subsequently lead to synapse loss. How excitatory circuit inputs are altered across the dendritic tree of individual neurons under neuroinflammatory stress is not well understood. Here, we directly assessed the structural dynamics of labeled excitatory synapses in experimental autoimmune encephalomyelitis (EAE) as a model of immune-mediated cortical neuronal damage. We used in vivo two-photon imaging and a synthetic tissue-hydrogel super-resolution imaging technique to reveal the dynamics of excitatory synapses, map their location across the dendritic tree of individual neurons, and examine neurons at super-resolution for synaptic loss. We found that excitatory synapses are destabilized but not lost from dendritic spines in EAE, starting with the earliest imaging session before symptom onset. This led to dramatic changes in excitatory circuit inputs to individual cells. In EAE, stable synapses are replaced by synapses that appear or disappear across the imaging sessions or repeatedly change at the same location. These unstable excitatory inputs occur closer to one another in EAE than in healthy controls and are distributed across the dendritic tree. When imaged at super-resolution, we found that a small proportion of dendritic protrusions lost their presynapse and/or postsynapse. Our finding of diffuse destabilizing effects of neuroinflammation on excitatory synapses across cortical neurons may have significant functional consequences since normal dendritic spine dynamics and clustering are essential for learning and memory.

## 1. Introduction

Although the pathologic hallmark of MS is demyelinating lesions in the central nervous system(Frischer et al., 2015), it is now recognized that another main pathology is neuronal loss(Carassiti et al., 2018) and damage with the enormous loss of synapses. This synapse loss is widespread, impacting excitatory and inhibitory synapses and involving the brain and spinal cord in demyelinated and normal-appearing grey matter (NAGM) (Huiskamp et al., 2022; Jürgens et al., 2016; Petrova et al., 2020; Vercellino et al., 2021; Zoupi et al., 2021). How synapse loss and dysfunction in MS contribute to physical neurologic disability and the cognitive dysfunction that impacts the majority of PwMS(Benedict et al., 2020) is unknown, but it may be an essential contributor(Filippo et al., 2018). Most post-mortem studies of synapse loss in people who had MS are from patients with longstanding disease(Möck et al., 2021), but it is likely that this synapse loss begins early in the disease(Bevan et al., 2018), possibly even before the onset of the first relapse symptoms(Azevedo et al., 2015; Bjornevik et al., 2020; Cen et al., 2023; Cortese et al., 2016), and is accelerated by inflammatory disease activity(Cagol et al., 2022). While the mechanisms of synapse loss remain poorly understood, it is not simply the result of demyelination and axonal transection(Jürgens et al., 2016); rather, inflammation shapes synapses even far from demyelinating lesions.

In a seminal work, a massive scale of synapse loss was revealed by reconstructing cortical pyramidal neurons labeled with Golgi-Cox staining from people who had longstanding MS(Jürgens et al., 2016). They found a dramatic loss of dendritic spines, impacting all locations of the sampled cortex. Dendritic spines are the anatomical site of most excitatory synapses(Kasthuri et al., 2015), so this finding translates to the massive loss of excitatory synapses. Numerous other studies have investigated the loss of synaptic puncta by staining either pre- or postsynaptic proteins in human post-mortem brain and spinal cord, and these confirm the loss of excitatory synapses(Möck et al., 2021). These post-mortem studies show the long-term consequences of MS on synapses but do not inform on the early impacts of neuroinflammation on synapses that subsequently lead to synapse loss. To answer this question, the EAE model can be used to model the early impacts of neuroinflammation on synapses. A study in the EAE model suggested that neuroinflammation destabilizes synapses(Yang et al., 2013). Short dendritic segments were in vivo two-photon imaged in mice with fluorescent labeling of layer V pyramidal neurons, and there was an increased turnover of dendritic spines. Dendritic spines were used as a structural surrogate of excitatory synapses, but it is unknown to what extent dendritic spines retain their excitatory synapse in EAE. To date, no studies have directly assessed the structural dynamics of labeled excitatory synapses in EAE, and there has been no comprehensive evaluation of how excitatory circuit inputs are altered across the dendritic tree of individual neurons.

Here, we used a fluorescent protein fused to an excitatory postsynaptic scaffolding protein to test how the remodeling of excitatory synapses in layer 2/3 cortical cells are altered in EAE and show that excitatory synapses are destabilized but not lost from dendritic spines in EAE. We used a synthetic tissue-hydrogel super-resolution imaging technique, Magnified Analysis of Proteome (MAP)(Balcioglu et al., 2023; Ku et al., 2016; Park et al., 2021), to test if these unstable excitatory synapses retain their presynaptic partner. When imaged at super-resolution, we found that a small proportion of dendritic protrusions lost their presynapse and/or postsynapse. Our findings indicate that in the setting of neuroinflammation, dendritic protrusions become unstable, and a small proportion lose their excitatory synapse. We longitudinally imaged the dendritic tree of individual cells to provide a comprehensive view of how the instability of excitatory synapses impacts excitatory circuit inputs to individual neurons. We show dramatic changes in excitatory inputs to individual cells, with stable synapses being replaced by synapses that appear or disappear across the imaging sessions or repeatedly change at the same location. The spacing of stably integrated synaptic excitatory inputs does not change in EAE, but unstable inputs occur in much closer proximity to one another and to stable excitatory inputs. These unstable synaptic inputs were not targeted to specific locations on the dendritic tree but instead were distributed across the dendritic tree.

## 2. Methods

### 2.1. Animals

All animal procedures were conducted in accordance with protocols approved by the Massachusetts General Hospital Institutional Animal Care and Use Committee (IACUC), and animals were cared for according to the requirements of the National Research Council’s Guide for the Care and Use of Laboratory Animals. All mice were C57BL/6J strain (Jackson # 000664).

### 2.2. DNA constructs

Generation of the Cre plasmid, and Cre dependent eYFP plasmid (*pSin wPGK-Cre*; Addgene plasmid # 101242 and *pFUdioeYFPW*; Addgene plasmid # 73858, respectively) has been previously described(Chen et al., 2012; Subramanian et al., 2013). The Cre-dependent PSD95-Teal plasmid was a gift from Prof. Elly Nedivi at the Massachusetts Institute of Technology. PSD95-Teal was amplified from *FuPSD95TealW* (a gift from Jerry Chen) with the forward and reverse primers containing an added Nhe1 site and MluI (complementary to Asc1) site, respectively. This amplified product was used to replace eYFP between Asc1 and Nhe1 sites of *FudioeYFPW*.

### 2.3. Surgical procedures

In utero electroporation on timed pregnant females was used to label layer 2/3 cortical pyramidal neurons with Enhanced yellow fluorescent protein (EYFP) for cell fill and to label excitatory synapses with a fusion protein, PSD95 fused to teal fluorescent protein (mTFP1), as previously described(Balcioglu et al., 2023; Tabata and Nakajima, 2001; Villa et al., 2016). At embryonic day 15.5, the pregnant mouse was anesthetized with isoflurane, and the uterus was elevated out of the abdomen through a midline incision. Then 0.75 µl of a plasmid solution containing the Cre plasmid 0.015 µg/µl, the Cre-dependent eYFP plasmid 0.7 µg/µl, and the Cre-dependent PSD95-Teal plasmid 0.25 µg/µl was injected into the right lateral ventricle of each fetus with a 32-gauge Hamilton syringe and needle (Hamilton Company # 7803-04 and 7634-01). A platinum 5 mm tweezer electrode (BTX # 450489) was placed with the positive electrode over the right somatosensory cortex of the fetus, and five pulses were delivered at 35 V (duration 50 ms, frequency 1 Hz) from a square-wave electroporator (ECM 830, BTX # 45-0662). At 7-8 weeks of age, male and female mice resulting from the in utero electroporation underwent implantation of a 5 mm cranial window over the right somatosensory cortex, as previously described with minor modifications(Holtmaat et al., 2009). Under isoflurane anesthesia, a micro-drill was used to create a 5 mm diameter opening in the right parietal bone. After removing the skull, a 5 mm glass coverslip was secured over the opening with dental cement and glue. A head plate (Narishige # CP-2) was secured with dental cement for chronic in vivo imaging.

### 2.4. In vivo two-photon imaging

Longitudinal in vivo two-photon imaging started ∼3 weeks after cranial window surgery at 11-15 weeks of age. Mice were anesthetized with isoflurane and were secured in a frame (Narishige # MAG-1) under the objective of the two-photon microscope. Two-photon imaging was done on an Olympus FluoView FV1000MPE multiphoton laser-scanning system mounted on an Olympus BX61WI microscope with a 25x objective (Olympus, NA= 1.05). A Mai Tai Ti: sapphire or InSight X3 laser (Spectra-Physics) generated two-photon excitation at 915 nm with power selected for no more than 50 mW of output from the objective. A well-isolated neuron in the primary somatosensory cortex by coordinates and with suitable YFP and PSD95-Teal expression was selected for longitudinal imaging in each mouse. An imaging volume was obtained to encompass the dendritic tree of the target neuron in 800 x 800 format with a resolution of 212 nm/pixel and 0.9 µm z step size.

### 2.5. EAE induction and clinical scoring

EAE was induced by injection of 200 μg MOG_35–55_ peptide emulsified in complete Freund’s adjuvant (CFA) with 2 subcutaneous injections of 0.1 ml of emulsion to each flank (Hooke # EK-2110). Pertussis toxin was given as a 0.1 ml intraperitoneal injection at a dose of 50 ng or 80 ng, due to different lot potency, in phosphate-buffered saline (PBS) on day 0 and either day 1 or 2 after immunization (Hooke # EK-2110). Clinical scores were given as follows: 0 no clinical signs, 1 tail paralysis, 2 tail paralysis and hind limb weakness, 3 tail paralysis and bilateral hind limb paralysis, 4 tail paralysis, complete bilateral hind limb paralysis and partial front limb paralysis, and 5 moribund. Of note, one EAE mouse that was longitudinally in vivo imaged had mild clinical EAE weakness with limp tail and hind limb inhibition but then developed a head tilt and falls for which a clinical score of 2.5 was given.

### 2.6. Magnified Analysis of Proteome (MAP) immunostaining and imaging

3 EAE mice and 2 healthy mice (15 – 22 weeks of age, male and female mice) resulting from the in utero electroporation and with cranial windows were transcardially perfused with ice-chilled PBS followed by 4% Paraformaldehyde (PFA). The EAE mice were perfused at maximum EAE clinical severity with clinical scores of 3-3.5 (14-16 days post-immunization). Brains were extracted and post-fixed in 4% PFA for 1 day at 4°C, with one brain having an additional 3 hrs of fixation in 4% PFA at room temperature (RT). Next, brains were washed in PBS at 4°C and then RT each for 1 day. A razor blade was used to take an ∼2 mm horizontal section of the cortex at the former location of the cranial window.

Cortical sections were processed for MAP and immunostained as previously described(Balcioglu et al., 2023). The cortical section was incubated overnight at 4°C in MAP solution (36 µl of initiator stock (10% (wt/vol) VA-044 (Wako Chemicals) in ice-cooled deionized water) added to 12 ml MAP stock (30% (wt/vol) acrylamide (MilliporeSigma # A9099), 10% (wt/vol) sodium acrylate (MilliporeSigma # 408220), and 0.1% bisacrylamide (Bio-Rad Laboratories # 161-0142) in PBS)). The section was then placed in a chamber created by adhering two microscope slides together with Blu-Tack (Bostik), and the chamber was filled with MAP solution. The chamber was inserted into a vented bottle (Chemglass Life Sciences # CLS-1429, Duran # Z742278) and connected to nitrogen gas for 2.5 hours until gelation was complete. The bottle was submerged in a bead bath set at 50°C and resting at a low angle (approximately 15°). The tissue-gel hybrid was hydrated overnight at 37°C in PBS followed by storage at RT, and then sectioned on a vibratome to obtain a ∼200 µm section at the brain’s surface. A tiled image of the blood vessels on the surface of the tissue-gel hybrid was obtained (Zeiss Axio Imager Z2) and aligned with an image of the blood vessels under the cranial window obtained when the mouse was alive. The somatosensory cortex was identified by coordinates and trimmed from the tissue-gel hybrid. The trimmed sample was cleared first by incubation for 4 – 6 hrs at 37°C in 12 ml clearing solution (6% (wt/vol) sodium dodecyl sulfate (MilliporeSigma # 75746), 50 mM sodium sulfite (MilliporeSigma # S0505), and 0.02% (wt/vol) sodium azide (MilliporeSigma # S2002) in 0.1 M phosphate buffer at pH 7.4) followed by immersion in 17.5 ml of clearing solution pre-heated to ∼ 86°C in a 95°C bead bath for either 5 or 10 min. The sample was then placed in a washing solution at RT (PBST - 0.1% (wt/vol) Triton X-100 in 1x PBS with 0.02% (wt/vol) sodium azide) and washed twice for a total of 6 hrs at 37°C, and then stored at RT.

The cleared sample was immunostained by incubation with primary antibodies in 1 ml PBST at 37°C or RT for 3-4 days, followed by 3 washes with PBST over 24 hours, followed by incubation with secondary antibodies in 500 µl PBST at 37°C or RT for 1 day, followed by 3 washes with PBST over 24 hours. Primary antibodies were as follows: PSD95 (UC Davis/NIH NeuroMab Facility # 73-028), Bassoon (Synaptic Systems # 141 004 and 141 003), GFP (Thermo Fisher Scientific # A10262). Secondary antibodies were as follows: Goat Anti-Guinea pig Alexa Fluor 405 (abcam # ab175678), Goat anti-Mouse Alexa Fluor Plus 488 (Thermo Fisher Scientific # A32723), Donkey Anti-Chicken Alexa Fluor 488 (Jackson ImmunoResearch Laboratories # 703-545-155), Goat Anti-Chicken Alexa Fluor 555 (abcam # ab150174), Donkey Anti-Mouse Alexa Fluor 555 (abcam # ab150106), Donkey Anti-Rabbit Alexa Fluor 647 (Thermo Fisher Scientific # A31573).

The stained sample was expanded in dilute buffer (0.02x PBS), resulting in a final expansion of ∼3-4 times expansion in all directions(Balcioglu et al., 2023). Samples were imaged free-floating in a glass-bottom dish covered with coverslips to immobilize the sample on an Olympus FV3000 Inverted Laser Scanning Confocal Microscope with a water immersion 60x objective (Olympus, NA=1.2). Individual dendrites on labeled neurons were selected for imaging favoring dendrites located within a continuous XY plane. They were imaged with a resolution of ∼100 nm/pixel and 0.5 µm z step size. Low-resolution whole-cell images of the anti-GFP labeled channel were obtained by obtaining a z-stack with a 10x objective (Olympus, NA=0.4).

### 2.7. Data and statistical analysis

Two-photon images were processed using the ImageJ (NIH) smooth command to decrease noise. Images were scored manually to track dendritic protrusions and whether they contained PSD95 with a modified version of the ObjectJ 4D point tracking system plugin for Fiji/ImageJ (https://sils.fnwi.uva.nl/bcb/objectj/index.html)(Villa et al., 2016). Dendritic protrusions were scored as previously described(Holtmaat et al., 2009), and protrusions in the z-plane were excluded from the analysis. For each scored cell, the researcher was blinded to the imaging session. A combined total of 3043 analyzed dendritic spines were present on the baseline imaging session for the 12 in vivo imaged cells. There was a mean of 254 ± 119 analyzed dendritic spines on the baseline imaging session for the in vivo imaged cells (Minimum 161, Maximum 510, Mean ± SD). Percent dynamic spines was calculated as ((N_gained_ + N_lost_)/ (N1_total_ + N2_total_)) X 100, as previously described(Subramanian et al., 2019) (N1 = total dendritic spines on the current session, N2 = total dendritic spines on next session). Percent spine loss was calculated as (N_lost_/ N1_total_) X 100, and percent spine gain was calculated as (N_gained_/ N2_total_) X 100. Spine density was calculated by tracing the scored dendritic segments in Neurolucida 360 (MBF Bioscience).

To examine the spatial organization of the spines across the dendritic tree, we reconstructed the dendritic trees of the neurons in Neurolucida 360 (MBF Bioscience). We manually placed makers at the location of the scored dendritic spines. Then, in Neurolucida Explorer, the Individual Marker Analysis was used to generate the distance of each dendritic spine along the branch. Analysis of nearest neighbor distance and spine density across the dendritic tree was done in Matlab. A Microsoft Excel file containing the distance of each scored dendritic spine from the cell body and along the scored dendritic segment was placed in a folder, the data path was specified, ensuring the Matlab code was on the path, and then the code was run. The code ‘Synapse_Analysis.mlx’ generates the nearest neighbor distance and then gives the synapse density /10 µm along each scored dendritic segment, with the bin advancing by 1 µm along the length of the segment for each column of the output.

MAP images were processed using the ImageJ (NIH) smooth command to decrease noise. Images were scored manually with a modified version of ObjectJ. Dendritic protrusions on the imaged dendrites were assessed for the presence of labeled PSD95 and/or Bassoon in the spine head.

GraphPad Prism version 9.4.1 for Windows (GraphPad Software) was used for statistical data analysis with t-test or ANOVA. IBM SPSS Statistics version 28.0 for Windows (IBM Corp.) was used for statistical data analysis with Chi-Square.

## 3. Results

### 3.1. Dendritic spines are unstable in EAE

To determine how neuroinflammation in the EAE model alters excitatory circuit inputs across individual cortical neurons, we used a strategy to label layer 2/3 neurons fluorescently and their complement of excitatory synapses. Then, we imaged these neurons using in vivo two-photon imaging longitudinally after induction of EAE. We used MOG_35-55_ EAE in C57BL/6 mice to model immune-mediated cortical neuronal damage. In the cerebral cortex, MOG_35-55_ EAE mice have cortical atrophy, neuron, axon, and synapse loss, microglia, and astrocyte activation, subpial demyelination, and small cortical demyelinating lesions(Burns et al., 2014; Errede et al., 2012; Girolamo et al., 2011; Hamilton et al., 2019). Plasmids were introduced into newborn neurons in fetal mice through in utero electroporation for labeling of neuronal cell fill with EYFP and labeling of the excitatory post-synaptic scaffolding protein PSD95 with a fusion protein, PSD95 fused to teal fluorescent protein (mTFP1), as previously described(Balcioglu et al., 2023; Villa et al., 2016)(Fig. 1a-b). After these mice were born and grew to young adulthood, a cranial window was implanted for in vivo two-photon imaging of well-isolated labeled neurons in the primary somatosensory cortex. Individual neurons were imaged weekly in healthy littermates, mice after induction of MOG_35-55_ EAE, and control mice that received injections of complete Freund’s adjuvant (CFA) and pertussis toxin but no MOG peptide. The fate of individual dendritic spines, the structural location of excitatory synapses, and whether they contained PSD95 was tracked across imaging session (Fig. 1c).

**Fig. 1.**
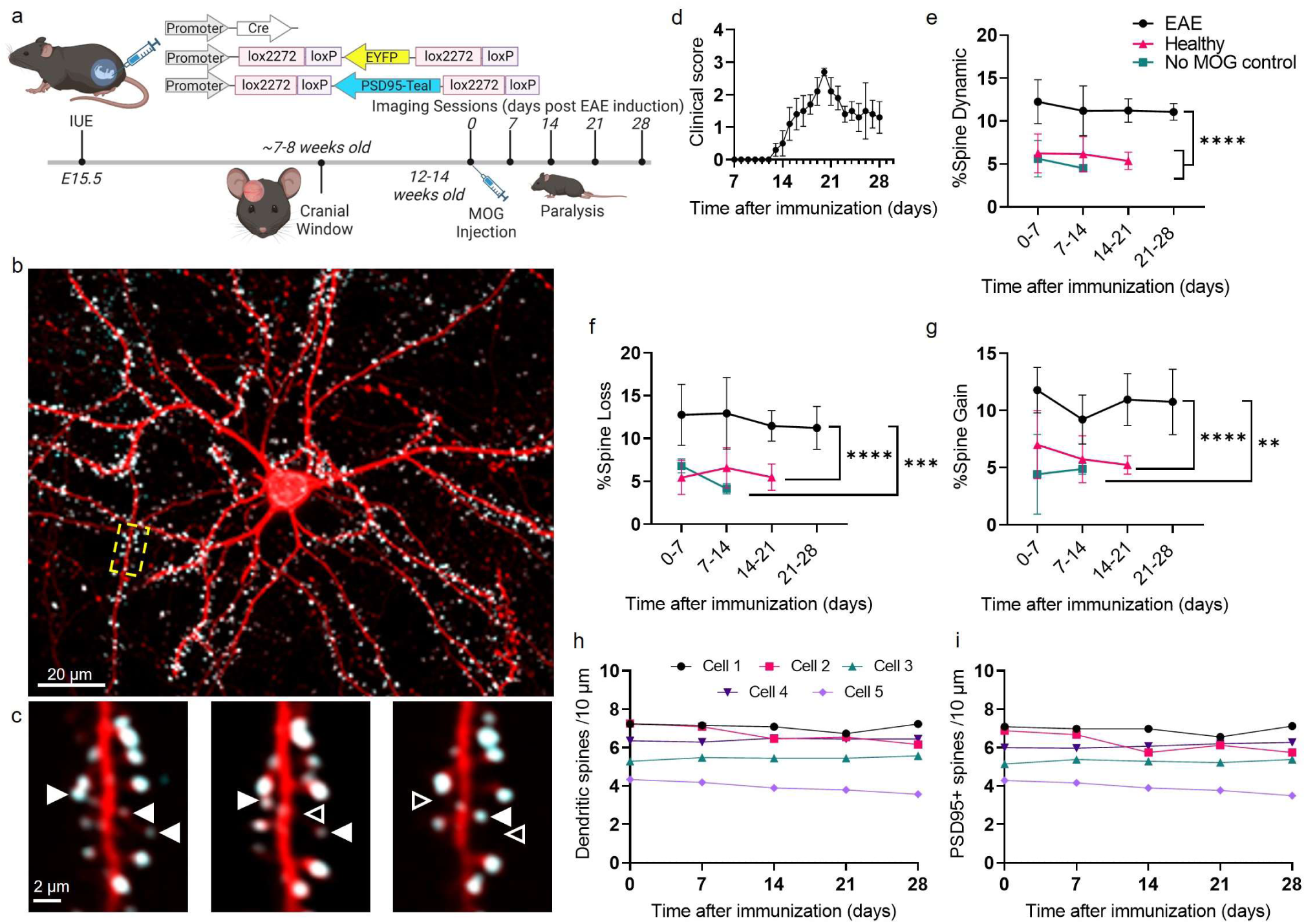
Excitatory synapses have increased structural dynamics after induction of EAE, but gains and losses are balanced, and there is no loss of overall dendritic spines or PSD95+ dendritic spines. **a** Experimental timeline and plasmids used for in utero electroporation and labeling of neuron cell fill (EYFP) and excitatory synapses (PSD95-Teal), created with BioRender.com. **b** Maximum intensity z-projection of layer 2/3 pyramidal cell imaged in vivo with YFP cell fill (red) and PSD95-Teal excitatory synapses (teal). Scale bar, 20 µm. **c** Magnified view of one x-y plane from the yellow-boxed dendritic segment from three consecutive weekly imaging sessions. Three dynamic excitatory synapses are marked with white triangles, filled triangles when the synapse is present on the imaging session, and open triangles when the synapse is absent on the imaging session. Scale bar, 2 µm. **d** EAE clinical scores for 5 mice imaged after induction of EAE. Mean ± SEM. **e** Percent dynamic dendritic spines (gains and losses) for mice subjected to EAE and imaged weekly until 28 days after immunization, healthy mice imaged weekly for 4 sessions, and no MOG control mice imaged weekly until 14 days after injection of CFA and pertussis toxin. Mean ± SD. **f, g** Percent dendritic spines lost (f) or gained (g). Mean ± SD. (n= 5 cells from 5 different EAE mice, 4 cells from 4 different healthy mice, 2 cells from 2 different no MOG control mice); **p = 0.0002 ***p = 0.0001 ****p <0.0001 (two-way ANOVA with Tukey’s post hoc test for multiple comparisons). See also Figure S2 for gains and losses in two EAE mice imaged at 6 weeks after induction of EAE. **h** Dendritic spine density, and **i** PSD95+ spine density after induction of EAE in 5 cells imaged from 5 different mice. p = 0.2866 for spine density, p = 0.3348 for PSD95+ spine density (Repeated measures ANOVA).

We imaged four healthy littermates for 4 weekly imaging sessions, and five mice subjected to MOG_35-55_ EAE and imaged at 0, 7, 14, 21, and 28 days after immunization. We also imaged control mice at 0, 7, and 14 days after injecting CFA and pertussis toxin. One additional mouse had two weekly baseline imaging sessions before immunization for MOG_35-55_ EAE and was imaged weekly until 14 days after immunization and there was a trend for increased mean weekly dendritic spine dynamics (gains and losses) in the 2 weeks after induction of EAE (Supplementary Fig. 1). The dynamics of dendritic spines were similar for healthy mice and control mice, with overall mean dendritic spine dynamics across all sessions of 5.9 ± 0.5 for healthy and 5.1 ± 0.8 for control (Fig. 1e, mean ± SD). Spine dynamics were significantly increased in EAE mice at 11.4 ± 0.5 (Fig. 1e, *F*(2, 30) = 37.58, p <0.0001, mean ± SD). Spine dynamics were increased in EAE mice before symptom onset at the earliest imaging session, comparing 0 to 7 days after immunization (Fig. 1e). They remained elevated through day 28 with no significant change over time. Gains and losses of spines were balanced (Fig. 1f,g). There was no significant decrease in the density of dendritic spines in EAE mice through day 28 (Fig. 1h). Since there was no decrease in spine density, these results indicate instability of excitatory inputs to cortical neurons rather than a large loss of excitatory inputs.

In two of the EAE mice, an additional imaging session was done at 6 weeks after immunization to determine if spine dynamics remained elevated at later time points. We found that dendritic spine gains remained elevated compared to healthy mice from day 28 to 42 after immunization (Supplementary Fig. 2b, one-tailed t-test, *t*(4) = 2.329, p = 0.0402), but spine losses were not significantly different. Of note, one of the EAE mice developed relapse symptoms before the final imaging session (Supplementary Fig. 2a). However, both mice had similar values for spine gains (11.4% and 10.1%), indicating that the elevated spine gains in the EAE mice from day 28 to 42 were not due to the relapse. This indicates that spine dynamics do not return to normal later in EAE.

### 3.2. Most dendritic spines in EAE contain excitatory synapses

We observed instability of dendritic spines in EAE, which led us to question if these unstable dendritic spines lose their excitatory synapse. On the baseline imaging session, before immunization for MOG_35-55_ EAE, 96.7% ± 2.0 of dendritic spines contained the post-synaptic protein PSD95 (Mean ± SD). There was no significant decrease in the density of dendritic spines containing PSD95 in EAE mice through day 28 (Fig. 1i). This indicates that dendritic spines retain an excitatory post-synaptic density in EAE. Still, the possibility remained that these excitatory synapses may have lost their presynaptic bouton partner.

To determine if excitatory synapses in cortical neurons in EAE retain the presynaptic compartment, we utilized MAP(Balcioglu et al., 2023; Ku et al., 2016; Park et al., 2021), a synthetic tissue-hydrogel super-resolution imaging technique. We examined synapses across the dendritic tree of somatosensory layer 2/3 cortical neurons from EAE mice with YFP and PSD95-Teal labeled neurons with brain tissue collection at peak clinical EAE severity. After processing the tissue for MAP, we immunostained for GFP to visualize cell fill, PSD95 to label the postsynapse, and Bassoon to label the presynapse (Fig. 2a-c). Grossly, synapses in EAE retained a normal appearance with a mix of small punctate synapses, horseshoe-shaped synapses, and perforated synapses (Fig. 2c). Most dendritic spines in the EAE cells contained an excitatory synapse with both presynaptic Bassoon and postsynaptic PSD95 (92.8%, Fig. 2d). However, this was significantly decreased from the healthy cells (97.7%, *X*^2^ (3, N = 1757) = 24.7, p <0.05)). Very few dendritic protrusions in healthy mice lacked an excitatory synapse (1.7%), and even fewer contained only PSD95 or Bassoon (0.2% and 0.4%, respectively, Fig. 2d). In EAE cells, there was a significant increase in dendritic protrusions that lacked an excitatory synapse (4%), had only PSD95 (1.7%) or only Bassoon (1.4%, *X*^2^ (3, N = 1757) = 24.7, p <0.05)). Due to the ability of MAP to image larger volumes at super-resolution, we obtained a large sample size of synapses, including very thin protrusions that cannot be imaged at the lower resolution of our in vivo two-photon experiments. These technical advantages allowed us to detect this loss of normal excitatory synapses in a small proportion of dendritic protrusions in EAE, indicating that in the setting of neuroinflammation, dendritic protrusions become unstable and a small proportion lose their excitatory synapse.

**Fig. 2.**
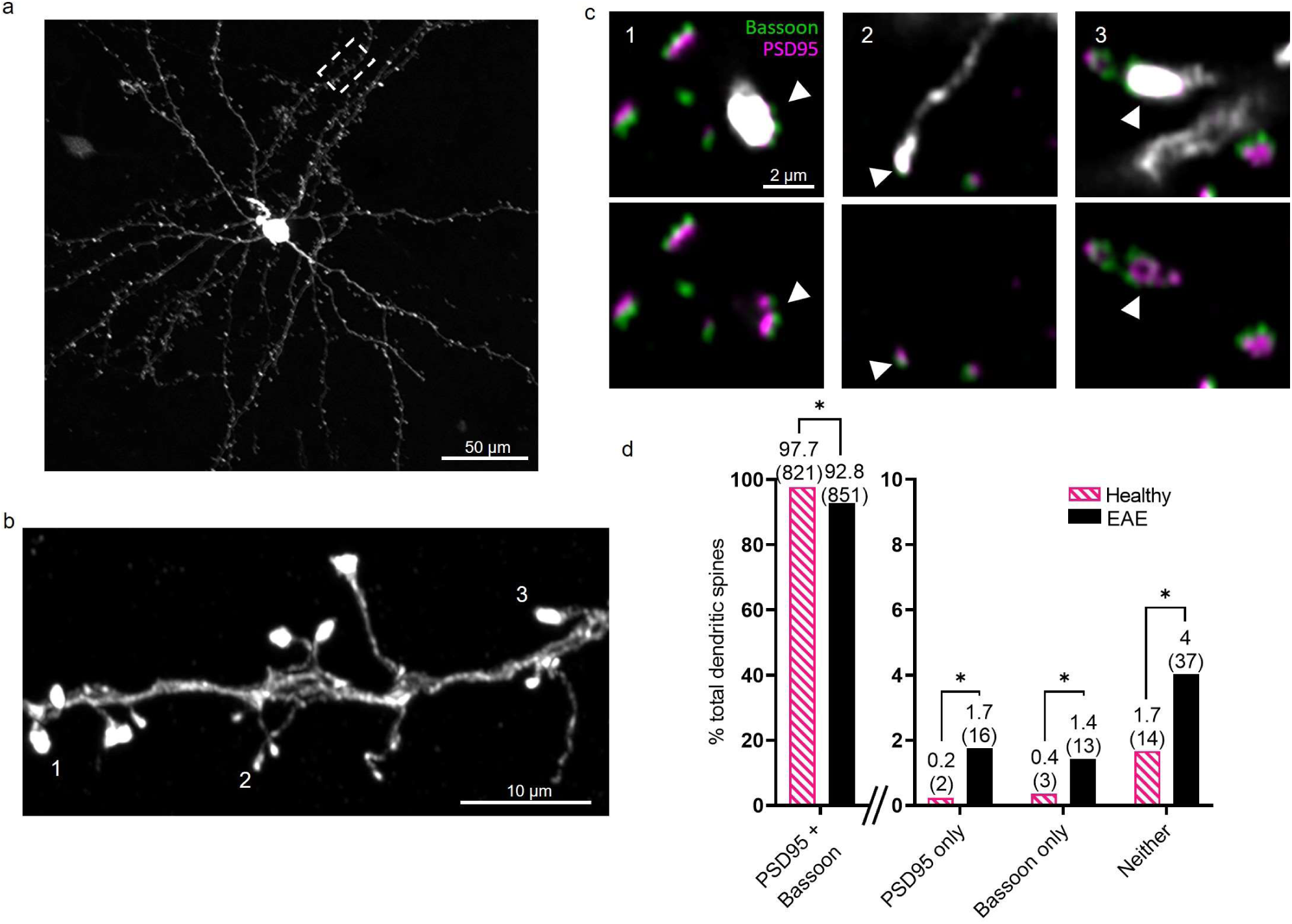
A small proportion of dendritic spines in EAE lose both presynaptic (bassoon) and postsynaptic (PSD95) scaffolding proteins, although most dendritic spines retain normal synaptic structure with the presence of both PSD95 and Bassoon. **a** Maximum intensity z-projection of layer 2/3 pyramidal cell in cleared and expanded brain tissue from a mouse with YFP and PSD95-Teal labeled neurons. Cell fill, anti-GFP. Scale bar, 50 µm. **b** Magnified image of one x-y plane from the white-boxed dendritic segment. Scale bar, 10 µm. **c** Magnified view of 3 dendritic spines numbered in b. The top shows cell fill (anti-GFP, white), anti-PSD95 (magenta), and anti-Bassoon (green), and the bottom shows only anti- PSD95 and anti-Bassoon. The location of the synapse on the dendritic spine head is marked with a triangle. Scale bar, 2 µm. **d** Percent and the number of dendritic spines that contain PSD95 and/or Bassoon, or neither in 3 cells from 3 different EAE mice at peak EAE scores of 3-3.5 (14-16 days post- immunization), and 2 cells from 2 different healthy mice. *p <0.05 (Chi-Square with Bonferroni correction for multiple comparisons).

### 3.3. Classifying dendritic spines by their dynamic behavior in EAE

Given the increased dendritic spine dynamics in EAE, we next examined how the dynamic behavior of spines is altered across cortical neurons in EAE. We classified dendritic spines by their dynamic behavior across four weekly imaging sessions at 7, 14, 21, and 28 days after immunization for the 5 EAE mice, and across four weekly imaging sessions for the 4 healthy mice. We defined a spine that was present on all 4 imaging sessions as stable, that has a one-time change (appear or disappear) as a one-time dynamic, that is gained and then lost as transient, and that is lost and regained at the exact location at least once as recurrent (Fig. 3a), like as in (Villa et al., 2016). As expected, in EAE, the proportion of stable spines was significantly decreased (61.9%) from healthy (75.7%, Fig. 3b, Supplementary Fig.3, *X*^2^ (3, N = 2771) = 70.756, p <0.05)). The stable spines lost in EAE were replaced by one-time dynamic and recurrent spines, and the proportion of recurrent spines more than doubled from 2.7% in healthy to 7.6% in EAE. This indicates that while dendritic spines are destabilized in EAE, the neurons retain the capacity to make new excitatory synaptic connections. In addition, they can reestablish lost connections, a behavior that is rare in healthy neurons.

**Fig. 3.**
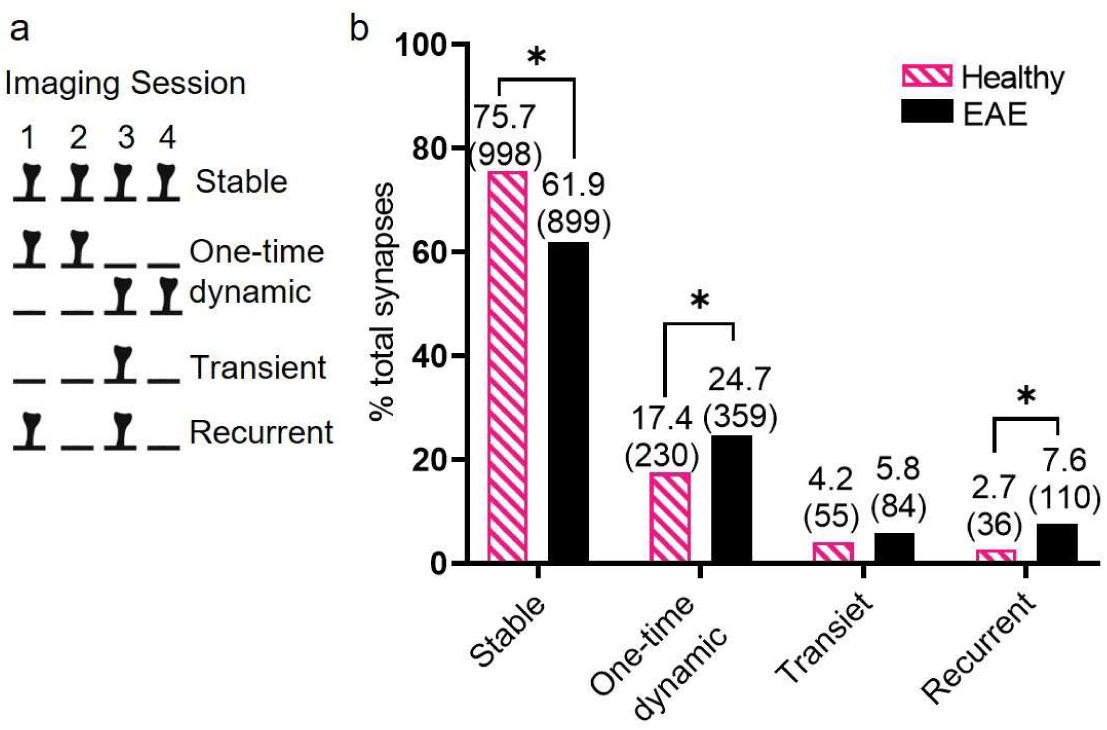
Classifying dendritic spines by their dynamic behavior in EAE revealed that stable spines are replaced in EAE by those that change one time during the imaging sessions (appear or disappear) and those that change more than one time at the same location. **a** Dendritic spine behavior was classified across four weekly imaging sessions at 7, 14, 21, and 28 days after immunization for the 5 EAE mice and across four weekly imaging sessions for the 4 healthy mice. **b** Percent and the number of dendritic spines in each category. See also Figure S3 for the proportion of spine types for each individual cell. *p<0.05 (Chi-Square with Bonferroni correction for multiple comparisons).

### 3.4. Distribution of dynamic and stable synapses on the dendritic tree in EAE

We next examined the spatial organization of the dynamic spines across the dendritic tree. We reconstructed the 5 EAE cells and 4 healthy cells and placed markers at the location of stable and dynamic dendritic spines (one-time dynamic, transient and recurrent combined) to generate the location of each spine relative to other spines on the branch and the cell body (Fig. 4a). We calculated the nearest neighbor distance for dynamic dendritic spines to the closest dynamic or stable spine, and for stable dendritic spines to the closest dynamic or stable spine. The average distance for dynamic-to-dynamic spines was cut in half in EAE compared to healthy (4.5 µm ± 1.5 for healthy and 2 µm ± 0.8 for EAE, Fig. 4b, *F*(1, 28) = 17.62, p = 0.0001, mean ± SD). The average distance for stable-to-dynamic was also cut in half in EAE compared to healthy (4 µm ± 0.9 for healthy and 2.1 µm ± 0.8 for EAE, p = 0.0027, mean ± SD). There was no difference in average distance for dynamic-to-stable or stable-to-stable. This indicates that while the spacing of stably integrated excitatory inputs does not change in EAE, dynamic inputs in much closer proximity will influence these stable inputs.

**Fig. 4.**
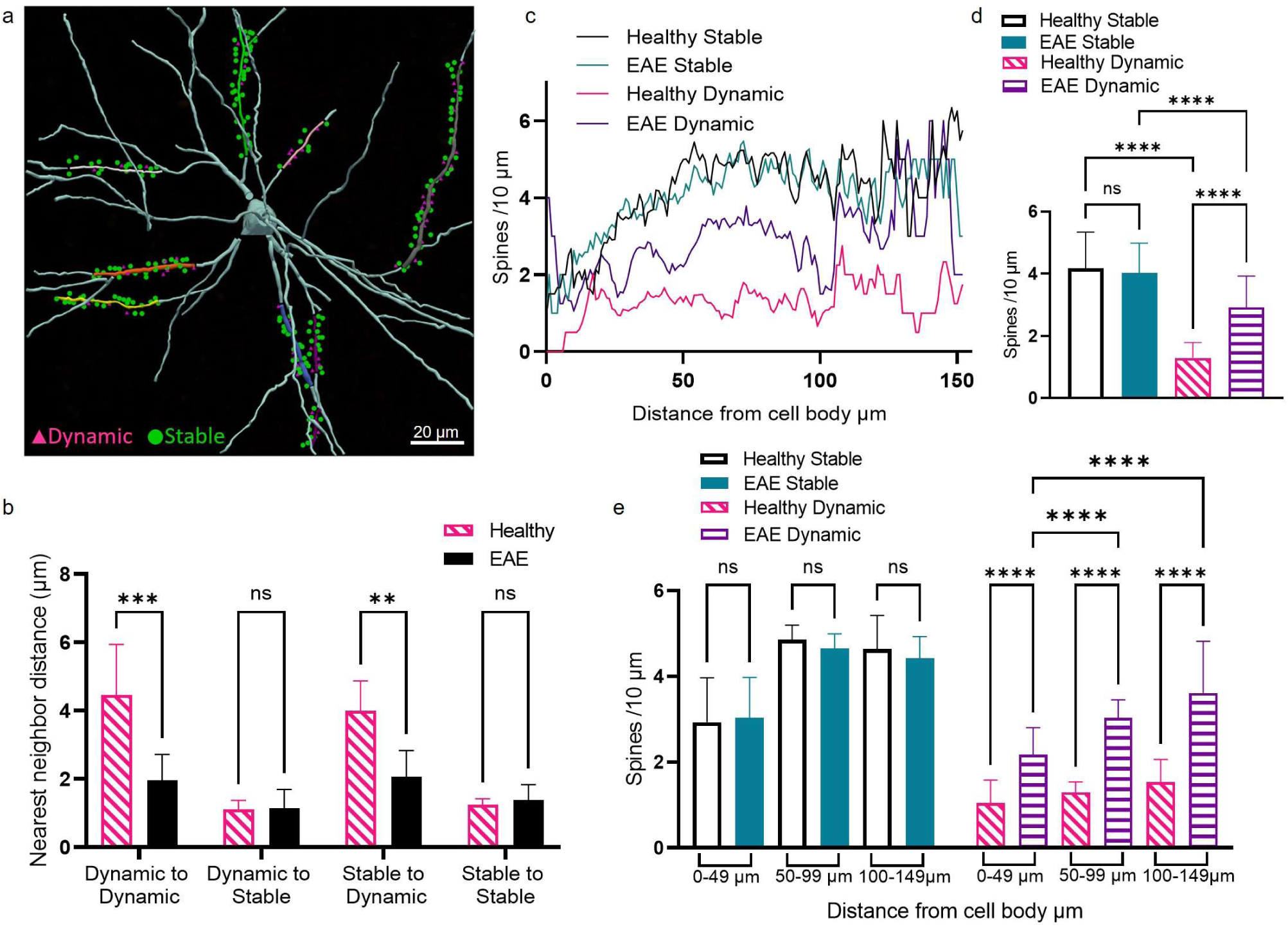
Dynamic dendritic spines are nearer to one another, and they are increased in density across the dendritic tree in EAE, with the most significant increase in density in the more distal dendritic tree >50 µm from the cell body. **a** The 5 EAE cells and 4 healthy cells were reconstructed in Neurolucida 360, and markers were placed for the location of dendritic spines categorized as depicted in Fig. 3 as stable or dynamic (one-time dynamic, transient, or recurrent). Cell body and non-analyzed branches = Cyan. **b** Nearest neighbor distance for dynamic dendritic spines to the closest dynamic or stable spine, and for stable dendritic spines to the closest dynamic or stable spine. Mean ± SD. **p = 0.0027 ***p = 0.0001 (two-way ANOVA with Šidák’s post hoc test for multiple comparisons). **c** Mean density of dynamic and stable spines for EAE and healthy cells by distance from the cell body. See also Figure S4 for graphs for each individual cell. **d** Mean overall density of dynamic and stable spines for EAE and healthy cells. Mean ± SD. ****p <0.0001 (two-way ANOVA with Tukey’s post hoc test for multiple comparisons). **e** Mean density of dynamic and stable spines for EAE and healthy cells for 0-49 µm from the cell body, 50-99 µm from the cell body, and 100-149 µm from the cell body. Mean ± SD. ****p <0.0001 (two-way ANOVA with Tukey’s post hoc test for multiple comparisons).

We next aimed to determine if these increased dynamic dendritic spines were targeted to specific locations in the dendritic tree. We calculated the density of dynamic dendritic spines and stable dendritic spines relative to the distance from the cell body (Fig. 4c, Supplementary Fig. 4). As expected, we found that the density of dynamic spines was significantly increased in EAE compared to healthy (2.9 spines/10 µm ± 1.0 for EAE and 1.3 ± 0.5 for healthy, Fig. 4d, *F*(3, 1420) = 426.6, p <0.0001, mean ± SD). We noticed the density of dynamic spines in EAE was lower proximal to the cell body and increased with increasing distance from the cell body (Fig. 4c). To examine this observation further; we separated the density of stable and dynamic spines into bins of 0-49 µm, 50-99 µm and 100-149 µm from the cell body (Fig. 4e). We found that dynamic spine density in EAE are significantly increased in the 50-99 µm and 100-149 µm bins compared to the 0-49 µm bin (*F*(11, 1959) = 221.6, p <0.0001). A similar trend was present for dynamic spines in healthy, where dynamic spine density was increased in the 100-149 µm bin compared to the 0-49 µm bin (p = 0.0441), but there was no difference between the 0-49 µm and 50-99 µm bins. Stable spines showed a similar pattern with lower density proximal to the cell body and increased density with increasing distance from the cell body (Fig. 4c,e), but there was no difference between EAE and healthy. This indicates that increased dynamic spines in EAE are not targeted to specific locations on the dendritic tree but develop in a normal distribution across the dendritic tree, mirroring the proximal to the distal increased density pattern observed in healthy neurons.

## 4. Discussion

Despite the use of highly effective disease modifying therapies (DMT) targeting the immune mechanisms of MS, many patients still develop cognitive dysfunction(Landmeyer et al., 2020) and worsening physical neurologic disability(Kappos et al., 2020; Wolinsky et al., 2020). A barrier to developing novel therapeutics to prevent this cognitive dysfunction and worsening neurologic disability is an incomplete understanding of the neurobiology of MS. Specifically; we have limited knowledge about how different populations of neurons and their synapses are dynamically perturbed by neuroinflammation in MS.

By longitudinally imaging excitatory synapses across the dendritic tree of individual cortical neurons during the course of neuroinflammation in EAE, we obtained a comprehensive view of how excitatory circuit inputs to cortical neurons are dynamically altered in the setting of neuroinflammation. We found that excitatory synapses are destabilized in EAE, dramatically impacting excitatory circuit inputs to individual neurons. Unstable synapses replace stably integrated synaptic excitatory inputs, and these unstable synapses become much closer in proximity to one another on the dendritic tree. While excitatory synapses are destabilized, this does not translate into a widespread loss of excitatory synapses. When examined at super-resolution, most dendritic protrusions retained an excitatory synapse, although there was a small proportion of dendritic protrusions that lost their synapse in EAE.

Mapping of unstable excitatory synapses in EAE onto the dendritic tree of layer 2/3 pyramidal cells revealed that they are distributed across the dendritic tree and are not targeted to specific areas of the dendritic tree. This supports that neuroinflammation in EAE has diffuse destabilizing effects on excitatory synapses across cortical neurons, which is likely to have important functional consequences. Dendritic spine turnover in the healthy brain is normal and critical for learning and memory(Runge et al., 2020). The turnover of dendritic spines in the healthy brain is important for the clustering of new dendritic spines to support learning and memory(Frank et al., 2018). The formation of new dendritic spines in the healthy brain is not random but is targeted to clusters of co-active dendritic spines in the setting of learning(Hedrick et al., 2022). In the setting of EAE, where excitatory synapses are diffusely destabilized, it is likely that normal spine formation and clustering to support learning and memory are altered. One study supports this hypothesis, where in a spontaneous model of EAE using transgenic mice that express myelin basic protein-specific T cell receptors, there were altered spine dynamics in response to training on a motor task before the onset of clinical symptoms(Huang et al., 2021). After motor training, both control and EAE mice had higher gains than losses, but while control mice added spines in response to motor training, the EAE mice did not lose as many preexisting spines. This may mean that control mice could target new dendritic spines to dendritic spine clusters to support learning. In contrast, the spontaneous EAE mice maintained preexisting spines that were not necessarily clustered.

Transcranial magnetic stimulation (TMS) studies in PwMS support that alterations in cortical plasticity observed in EAE may have relevance to human disease. With a motor task, PwMS have impaired cortical plasticity in the motor cortex(Bassi et al., 2020), and cortical plasticity was reduced in PwMS with cognitive dysfunction(Balloff et al., 2022). This suggests that PwMS have alterations in cortical plasticity to learning and that this may be clinically meaningful for MS symptoms. The use of PET ligands targeting synapses with synaptic vesicle protein 2A (SV2A) ligands may, for the first time, reveal dynamic alterations in synapses during the course of MS(Carson et al., 2022), and resolve questions about when synapse loss begins in PwMS. However, SV2A PET measures synaptic density, and as demonstrated in EAE models, alterations in synapse structural stability, even without gross loss of synapses, can have profound effects on cortical connectivity.

There is an incomplete understanding of the mechanisms that lead to diffuse synaptic dysfunction and loss in MS, even far from inflammatory demyelinating lesions. However, it is likely to be a multifactorial process. Possible contributors include soluble proinflammatory molecules, neurotoxic astrocytes, synapse-engulfing microglia, and mitochondrial dysfunction(Bellingacci et al., 2021). Inflammatory pathways that have been implicated include complement (C1q and C3)(Bourel et al., 2021; Hammond et al., 2020; Michailidou et al., 2015; Ramaglia et al., 2021; Werneburg et al., 2020), Interleukin-1β (IL-1β) and tumor necrosis factor α (TNFα) (Rizzo et al., 2018; Yang et al., 2013). Astrocytes physically touch synapses and regulate synaptic function(Chung et al., 2015), and astrocyte signaling leads to synaptic dysfunction in EAE(Habbas et al., 2015). Microglia have been shown to engulf synapses, both in acute demyelinating lesions(Jafari et al., 2021), and in grey matter unaffected by demyelination(Werneburg et al., 2020). Mitochondrial dysfunction may lead to inadequate energy supplies to support synaptic function(Faria-Pereira and Morais, 2022; Sadeghian et al., 2016). The multiple possible mechanisms for synaptic dysfunction and loss in MS represent opportunities for possible therapeutic targets for neuroprotection in PwMS.

In summary, we found that excitatory synapses are destabilized in EAE, which may dramatically impact the integration of excitatory synaptic inputs to individual neurons. We found that despite increased dendritic spine dynamics in EAE, most dendritic protrusions retain a synapse. This suggests that, given the correct conditions, normal excitation can be restored to cortical neurons impacted by neuroinflammation.

## Acknowledgments

We thank B. Trippe for technical support. We thank M. Algamal for comments on the manuscript. This work was supported by NIH K08NS107591 (RG), and NIH R56AG060974 (BB).

## Notes

### Competing Interest Statement

The authors have declared no competing interest.

## References Cited

Azevedo, C.J., Overton, E., Khadka, S., Buckley, J., Liu, S., Sampat, M., Kantarci, O., Frenay, C.L., Siva, A., Okuda, D.T., Pelletier, D., 2015. Early CNS neurodegeneration in radiologically isolated syndrome. Neurology - Neuroimmunol Neuroinflammation 2, e102. 10.1212/nxi.0000000000000102

Balcioglu, A., Gillani, R., Doron, M., Burnell, K., Ku, T., Erisir, A., Chung, K., Segev, I., Nedivi, E., 2023. Mapping thalamic innervation to individual L2/3 pyramidal neurons and modeling their ‘readout’ of visual input. Nat. Neurosci. 26, 470–480. 10.1038/s41593-022-01253-9

Balloff, C., Penner, I.-K., Ma, M., Georgiades, I., Scala, L., Troullinakis, N., Graf, J., Kremer, D., Aktas, O., Hartung, H.-P., Meuth, S.G., Schnitzler, A., Groiss, S.J., Albrecht, P., 2022. The degree of cortical plasticity correlates with cognitive performance in patients with Multiple Sclerosis. Brain Stimul. 15, 403–413. 10.1016/j.brs.2022.02.007

Bassi, M.S., Buttari, F., Maffei, P., Paolis, N.D., Sancesario, A., Gilio, L., Pavone, L., Pasqua, G., Simonelli, I., Sica, F., Fantozzi, R., Bellantonio, P., Centonze, D., Iezzi, E., 2020. Practice-dependent motor cortex plasticity is reduced in non-disabled multiple sclerosis patients. Clin. Neurophysiol. 131, 566–573. 10.1016/j.clinph.2019.10.023

Bellingacci, L., Mancini, A., Gaetani, L., Tozzi, A., Parnetti, L., Filippo, M.D., 2021. Synaptic Dysfunction in Multiple Sclerosis: A Red Thread from Inflammation to Network Disconnection. Int J Mol Sci 22, 9753. 10.3390/ijms22189753

Benedict, R.H.B., Amato, M.P., DeLuca, J., Geurts, J.J.G., 2020. Cognitive impairment in multiple sclerosis: clinical management, MRI, and therapeutic avenues. Lancet Neurol. 19, 860–871. 10.1016/s1474-4422(20)30277-5

Bevan, R.J., Evans, R., Griffiths, L., Watkins, L.M., Rees, M.I., Magliozzi, R., Allen, I., McDonnell, G., Kee, R., Naughton, M., Fitzgerald, D.C., Reynolds, R., Neal, J.W., Howell, O.W., 2018. Meningeal inflammation and cortical demyelination in acute multiple sclerosis. Ann Neurol 84, 829–842. 10.1002/ana.25365

Bjornevik, K., Munger, K.L., Cortese, M., Barro, C., Healy, B.C., Niebuhr, D.W., Scher, A.I., Kuhle, J., Ascherio, A., 2020. Serum Neurofilament Light Chain Levels in Patients With Presymptomatic Multiple Sclerosis. Jama Neurol 77, 58–64. 10.1001/jamaneurol.2019.3238

Bourel, J., Planche, V., Dubourdieu, N., Oliveira, A., Séré, A., Ducourneau, E.-G., Tible, M., Maitre, M., Lesté-Lasserre, T., Nadjar, A., Desmedt, A., Ciofi, P., Oliet, S.H., Panatier, A., Tourdias, T., 2021. Complement C3 mediates early hippocampal neurodegeneration and memory impairment in experimental multiple sclerosis. Neurobiol Dis 160, 105533. 10.1016/j.nbd.2021.105533

Burns, T., Miers, L., Xu, J., Man, A., Moreno, M., Pleasure, D., Bannerman, P., 2014. Neuronopathy in the Motor Neocortex in a Chronic Model of Multiple Sclerosis. J Neuropathology Exp Neurology 73, 335–344. 10.1097/nen.0000000000000058

Cagol, A., Schaedelin, S., Barakovic, M., Benkert, P., Todea, R.-A., Rahmanzadeh, R., Galbusera, R., Lu, P.-J., Weigel, M., Melie-Garcia, L., Ruberte, E., Siebenborn, N., Battaglini, M., Radue, E.-W., Yaldizli, Ö., Oechtering, J., Sinnecker, T., Lorscheider, J., Fischer-Barnicol, B., Müller, S., Achtnichts, L., Vehoff, J., Disanto, G., Findling, O., Chan, A., Salmen, A., Pot, C., Bridel, C., Zecca, C., Derfuss, T., Lieb, J.M., Remonda, L., Wagner, F., Vargas, M.I., Pasquier, R.D., Lalive, P.H., Pravatà, E., Weber, J., Cattin, P.C., Gobbi, C., Leppert, D., Kappos, L., Kuhle, J., Granziera, C., 2022. Association of Brain Atrophy With Disease Progression Independent of Relapse Activity in Patients With Relapsing Multiple Sclerosis. Jama Neurol 79, 682–692. 10.1001/jamaneurol.2022.1025

Carassiti, D., Altmann, D.R., Petrova, N., Pakkenberg, B., Scaravilli, F., Schmierer, K., 2018. Neuronal loss, demyelination and volume change in the multiple sclerosis neocortex. Neuropathol. Appl. Neurobiol. 44, 377–390. 10.1111/nan.12405

Carson, R.E., Naganawa, M., Toyonaga, T., Koohsari, S., Yang, Y., Chen, M.-K., Matuskey, D., Finnema, S.J., 2022. Imaging of Synaptic Density in Neurodegenerative Disorders. J. Nucl. Med. 63, 60S–67S. 10.2967/jnumed.121.263201

Cen, S., Gebregziabher, M., Moazami, S., Azevedo, C.J., Pelletier, D., 2023. Toward precision medicine using a “digital twin” approach: modeling the onset of disease-specific brain atrophy in individuals with multiple sclerosis. Sci. Rep. 13, 16279. 10.1038/s41598-023-43618-5

Chen, J.L., Villa, K.L., Cha, J.W., So, P.T., Kubota, Y., Nedivi, E., 2012. Clustered dynamics of inhibitory synapses and dendritic spines in the adult neocortex. Neuron 74, 361–73. 10.1016/j.neuron.2012.02.030

Chung, W.-S., Allen, N.J., Eroglu, C., 2015. Astrocytes Control Synapse Formation, Function, and Elimination. Csh Perspect Biol 7, a020370. 10.1101/cshperspect.a020370

Cortese, M., Riise, T., Bjørnevik, K., Bhan, A., Farbu, E., Grytten, N., Hogenesch, I., Midgard, R., Simonsen, C.S., Telstad, W., Ascherio, A., Myhr, K., 2016. Preclinical disease activity in multiple sclerosis: A prospective study of cognitive performance prior to first symptom. Ann Neurol 80, 616–624. 10.1002/ana.24769

Errede, M., Girolamo, F., Ferrara, G., Strippoli, M., Morando, S., Boldrin, V., Rizzi, M., Uccelli, A., Perris, R., Bendotti, C., Salmona, M., Roncali, L., Virgintino, D., 2012. Blood-Brain Barrier Alterations in the Cerebral Cortex in Experimental Autoimmune Encephalomyelitis. J Neuropathology Exp Neurology 71, 840–854. 10.1097/nen.0b013e31826ac110

Faria-Pereira, A., Morais, V.A., 2022. Synapses: The Brain’s Energy-Demanding Sites. Int. J. Mol. Sci. 23, 3627. 10.3390/ijms23073627

Filippo, M.D., Portaccio, E., Mancini, A., Calabresi, P., 2018. Multiple sclerosis and cognition: synaptic failure and network dysfunction. Nat Rev Neurosci 19, 599–609. 10.1038/s41583-018-0053-9

Frank, A., Huang, S., Zhou, M., Nature …, G.A., 2018. Hotspots of dendritic spine turnover facilitate clustered spine addition and learning and memory. 10.1038/s41467-017-02751-2

Frischer, J.M., Weigand, S.D., Guo, Y., Kale, N., Parisi, J.E., Pirko, I., Mandrekar, J., Bramow, S., Metz, I., Brück, W., Lassmann, H., Lucchinetti, C.F., 2015. Clinical and pathological insights into the dynamic nature of the white matter multiple sclerosis plaque. Ann. Neurol. 78, 710–721. 10.1002/ana.24497

Girolamo, F., Ferrara, G., Strippoli, M., Rizzi, M., Errede, M., Trojano, M., Perris, R., Roncali, L., Svelto, M., Mennini, T., Virgintino, D., 2011. Cerebral cortex demyelination and oligodendrocyte precursor response to experimental autoimmune encephalomyelitis. Neurobiol Dis 43, 678–689. 10.1016/j.nbd.2011.05.021

Habbas, S., Santello, M., Becker, D., Stubbe, H., Zappia, G., Liaudet, N., Klaus, F.R., Kollias, G., Fontana, A., Pryce, C.R., Suter, T., Volterra, A., 2015. Neuroinflammatory TNFα Impairs Memory via Astrocyte Signaling. Cell 163, 1730–1741. 10.1016/j.cell.2015.11.023

Hamilton, A.M., Forkert, N.D., Yang, R., Wu, Y., Rogers, J.A., Yong, V.W., Dunn, J.F., 2019. Central nervous system targeted autoimmunity causes regional atrophy: a 9.4T MRI study of the EAE mouse model of Multiple Sclerosis. Sci Rep-uk 9, 8488. 10.1038/s41598-019-44682-6

Hammond, J.W., Bellizzi, M.J., Ware, C., Qiu, W.Q., Saminathan, P., Li, H., Luo, S., Ma, S.A., Li, Y., Gelbard, H.A., 2020. Complement-dependent synapse loss and microgliosis in a mouse model of multiple sclerosis. Brain Behav Immun 87, 739–750. 10.1016/j.bbi.2020.03.004

Hedrick, N.G., Lu, Z., Bushong, E., Singhi, S., Nguyen, P., Magaña, Y., Jilani, S., Lim, B.K., Ellisman, M., Komiyama, T., 2022. Learning binds new inputs into functional synaptic clusters via spinogenesis. Nat. Neurosci. 25, 726–737. 10.1038/s41593-022-01086-6

Holtmaat, A., Bonhoeffer, T., Chow, D.K., Chuckowree, J., Paola, V.D., Hofer, S.B., Hübener, M., Keck, T., Knott, G., Lee, W.-C.A., Mostany, R., Mrsic-Flogel, T.D., Nedivi, E., Portera-Cailliau, C., Svoboda, K., Trachtenberg, J.T., Wilbrecht, L., 2009. Long-term, high-resolution imaging in the mouse neocortex through a chronic cranial window. Nat. Protoc. 4, 1128–1144. 10.1038/nprot.2009.89

Huang, L., Lafaille, J.J., Yang, G., 2021. Learning-dependent dendritic spine plasticity is impaired in spontaneous autoimmune encephalomyelitis. Dev Neurobiol 81, 736–745. 10.1002/dneu.22827

Huiskamp, M., Kiljan, S., Kulik, S., Witte, M.E., Jonkman, L.E., Bol, J.G., Schenk, G.J., Hulst, H.E., Tewarie, P., Schoonheim, M.M., Geurts, J.J., 2022. Inhibitory synaptic loss drives network changes in multiple sclerosis: An ex vivo to in silico translational study. Mult. Scler. (Houndmills, Basingstoke, Engl.) 28, 2010–2019. 10.1177/13524585221125381

Jafari, M., Schumacher, A.-M., Snaidero, N., Gavilanes, E.M., Neziraj, T., Kocsis-Jutka, V., Engels, D., Jürgens, T., Wagner, I., Weidinger, J., Schmidt, S.S., Beltrán, E., Hagan, N., Woodworth, L., Ofengeim, D., Gans, J., Wolf, F., Kreutzfeldt, M., Portugues, R., Merkler, D., Misgeld, T., Kerschensteiner, M., 2021. Phagocyte-mediated synapse removal in cortical neuroinflammation is promoted by local calcium accumulation. Nature Neuroscience 24, 355–367. 10.1038/s41593-020-00780-7

Jürgens, T., Jafari, M., Kreutzfeldt, M., Bahn, E., Brück, W., Kerschensteiner, M., Merkler, D., 2016. Reconstruction of single cortical projection neurons reveals primary spine loss in multiple sclerosis. Brain : a journal of neurology 139, 39–46. 10.1093/brain/awv353

Kappos, L., Wolinsky, J.S., Giovannoni, G., Arnold, D.L., Wang, Q., Bernasconi, C., Model, F., Koendgen, H., Manfrini, M., Belachew, S., Hauser, S.L., 2020. Contribution of Relapse-Independent Progression vs Relapse-Associated Worsening to Overall Confirmed Disability Accumulation in Typical Relapsing Multiple Sclerosis in a Pooled Analysis of 2 Randomized Clinical Trials. Jama Neurol 77, 1132–1140. 10.1001/jamaneurol.2020.1568

Kasthuri, N., Hayworth, K., Berger, D., Schalek, R., Conchello, J., Knowles-Barley, S., Lee, D., Vázquez-Reina, A., Kaynig, V., Jones, T., 2015. Saturated reconstruction of a volume of neocortex. Cell 162, 648–661.

Ku, T., Swaney, J., Park, J.-Y., Albanese, A., Murray, E., Cho, J., Park, Y.-G., Mangena, V., Chen, J., Chung, K., 2016. Multiplexed and scalable super-resolution imaging of three-dimensional protein localization in size-adjustable tissues. Nature Biotechnology 34, 973–981. 10.1038/nbt.3641

Landmeyer, N.C., Bürkner, P.-C., Wiendl, H., Ruck, T., Hartung, H.-P., Holling, H., Meuth, S.G., Johnen, A., 2020. Disease-modifying treatments and cognition in relapsing-remitting multiple sclerosis: A meta-analysis. Neurology 94, e2373–e2383. 10.1212/wnl.0000000000009522

Michailidou, I., Willems, J.G., Kooi, E.-J.J., Eden, C. van, Gold, S.M., Geurts, J.J., Baas, F., Huitinga, I., Ramaglia, V., 2015. Complement C1q-C3-associated synaptic changes in multiple sclerosis hippocampus. Annals of neurology 77, 1007–26. 10.1002/ana.24398

Möck, E.E.A., Honkonen, E., Airas, L., 2021. Synaptic Loss in Multiple Sclerosis: A Systematic Review of Human Post-mortem Studies. Front Neurol 12, 782599. 10.3389/fneur.2021.782599

Park, Joha, Khan, S., Yun, D.H., Ku, T., Villa, K.L., Lee, J.E., Zhang, Q., Park, Juhyuk, Feng, G., Nedivi, E., Chung, K., 2021. Epitope-preserving magnified analysis of proteome (eMAP). Sci Adv 7, eabf6589. 10.1126/sciadv.abf6589

Petrova, N., Nutma, E., Carassiti, D., Newman, J.R., Amor, S., Altmann, D.R., Baker, D., Schmierer, K., 2020. Synaptic Loss in Multiple Sclerosis Spinal Cord. Ann Neurol 88, 619–625. 10.1002/ana.25835

Ramaglia, V., Dubey, M., Malpede, A.M., Petersen, N., Vries, S.I. de, Ahmed, S.M., Lee, D.S., Schenk, G.J., Gold, S.M., Huitinga, I., Gommerman, J.L., Geurts, J.J., Kole, M.H., 2021. Complement-associated loss of CA2 inhibitory synapses in the demyelinated hippocampus impairs memory. Acta Neuropathol 1–25. 10.1007/s00401-021-02338-8

Rizzo, F.R., Musella, A., Vito, F.D., Fresegna, D., Bullitta, S., Vanni, V., Guadalupi, L., Bassi, M.S., Buttari, F., Mandolesi, G., Centonze, D., Gentile, A., 2018. Tumor Necrosis Factor and Interleukin-1 β Modulate Synaptic Plasticity during Neuroinflammation. Neural Plast 2018, 1–12. 10.1155/2018/8430123

Runge, K., Cardoso, C., Chevigny, A. de, 2020. Dendritic Spine Plasticity: Function and Mechanisms. Front. Synaptic Neurosci. 12, 36. 10.3389/fnsyn.2020.00036

Sadeghian, M., Mastrolia, V., Haddad, A.R., Mosley, A., Mullali, G., Schiza, D., Sajic, M., Hargreaves, I., Heales, S., Duchen, M.R., Smith, K.J., 2016. Mitochondrial dysfunction is an important cause of neurological deficits in an inflammatory model of multiple sclerosis. Sci. Rep. 6, 33249. 10.1038/srep33249

Subramanian, J., Dye, L., Morozov, A., 2013. Rap1 Signaling Prevents L-Type Calcium Channel-Dependent Neurotransmitter Release. J. Neurosci. 33, 7245–7252. 10.1523/jneurosci.5963-11.2013

Subramanian, J., Michel, K., Benoit, M., Nedivi, E., 2019. CPG15/Neuritin Mimics Experience in Selecting Excitatory Synapses for Stabilization by Facilitating PSD95 Recruitment. Cell Rep. 28, 1584–1595.e5. 10.1016/j.celrep.2019.07.012

Tabata, H., Nakajima, K., 2001. Efficient in utero gene transfer system to the developing mouse brain using electroporation: visualization of neuronal migration in the developing cortex. Neuroscience 103, 865–872. 10.1016/s0306-4522(01)00016-1

Vercellino, M., Marasciulo, S., Grifoni, S., Vallino-Costassa, E., Bosa, C., Pasanisi, M.B., Crociara, P., Casalone, C., Chiò, A., Giordana, M.T., Corona, C., Cavalla, P., 2021. Acute and chronic synaptic pathology in multiple sclerosis gray matter. Multiple sclerosis (Houndmills, Basingstoke, England) 13524585211022174. 10.1177/13524585211022174

Villa, K.L., Berry, K.P., Subramanian, J., Cha, J.W., Oh, W.C., Kwon, H.-B.B., Kubota, Y., So, P.T., Nedivi, E., 2016. Inhibitory Synapses Are Repeatedly Assembled and Removed at Persistent Sites In Vivo. Neuron 89, 756–69. 10.1016/j.neuron.2016.01.010

Werneburg, S., Jung, J., Kunjamma, R.B., Ha, S.-K.K., Luciano, N.J., Willis, C.M., Gao, G., Biscola, N.P., Havton, L.A., Crocker, S.J., Popko, B., Reich, D.S., Schafer, D.P., 2020. Targeted Complement Inhibition at Synapses Prevents Microglial Synaptic Engulfment and Synapse Loss in Demyelinating Disease. Immunity 52, 167–182.e7. 10.1016/j.immuni.2019.12.004

Wolinsky, J.S., Arnold, D.L., Brochet, B., Hartung, H.-P., Montalban, X., Naismith, R.T., Manfrini, M., Overell, J., Koendgen, H., Sauter, A., Bennett, I., Hubeaux, S., Kappos, L., Hauser, S.L., 2020. Long-term follow-up from the ORATORIO trial of ocrelizumab for primary progressive multiple sclerosis: a post-hoc analysis from the ongoing open-label extension of the randomised, placebo-controlled, phase 3 trial. Lancet Neurology 19, 998–1009. 10.1016/s1474-4422(20)30342-2

Yang, G., Parkhurst, C.N., Hayes, S., Gan, W.-B.B., 2013. Peripheral elevation of TNF-α leads to early synaptic abnormalities in the mouse somatosensory cortex in experimental autoimmune encephalomyelitis. Proceedings of the National Academy of Sciences of the United States of America 110, 10306–11. 10.1073/pnas.1222895110

Zoupi, L., Booker, S.A., Eigel, D., Werner, C., Kind, P.C., Spires-Jones, T.L., Newland, B., Williams, A.C., 2021. Selective vulnerability of inhibitory networks in multiple sclerosis. Acta Neuropathol 141, 415–429. 10.1007/s00401-020-02258-z

