## Supplementary Figures for "Instability of excitatory synapses in experimental autoimmune encephalomyelitis and the outcome for excitatory circuit inputs to individual cortical neurons"

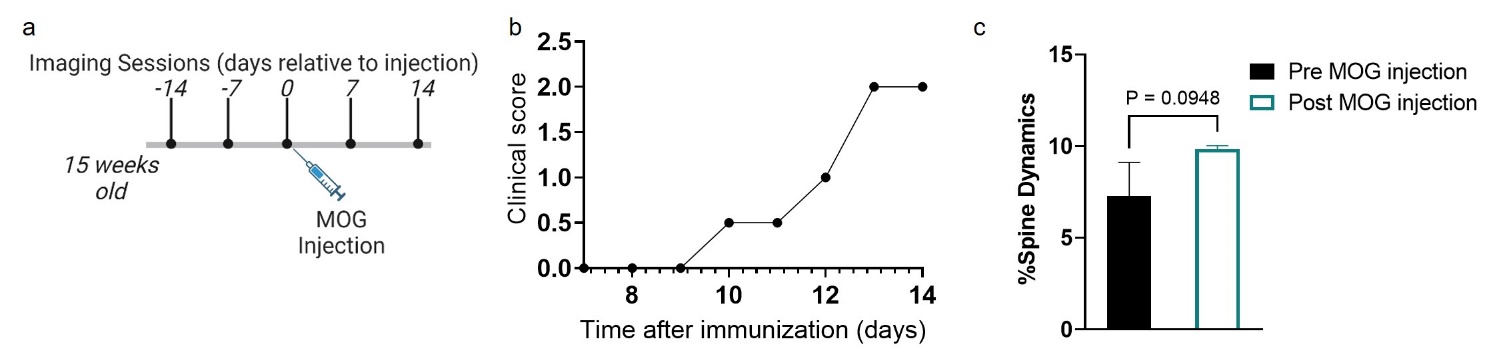

**Supp Fig. 1**

Mean weekly dendritic spine dynamics (gains and losses) in one mouse in the 2 weeks after induction of EAE, compared to the two baseline weeks of imaging. **a** Experimental timeline for imaging sessions. **b** EAE clinical scores for the imaged mouse. **C** Mean weekly dendritic spine dynamics before and after MOG injection showed a trend for an increase in the 2 weeks after induction of EAE 9.83 ± 0.20, compared to the two baseline weeks of imaging 7.30 ± 1.82. p = 0.0948. (one-tailed t-test, *t*(2) = 1.957). Mean ± SD. (n = 1 cells from 1 EAE mouse).

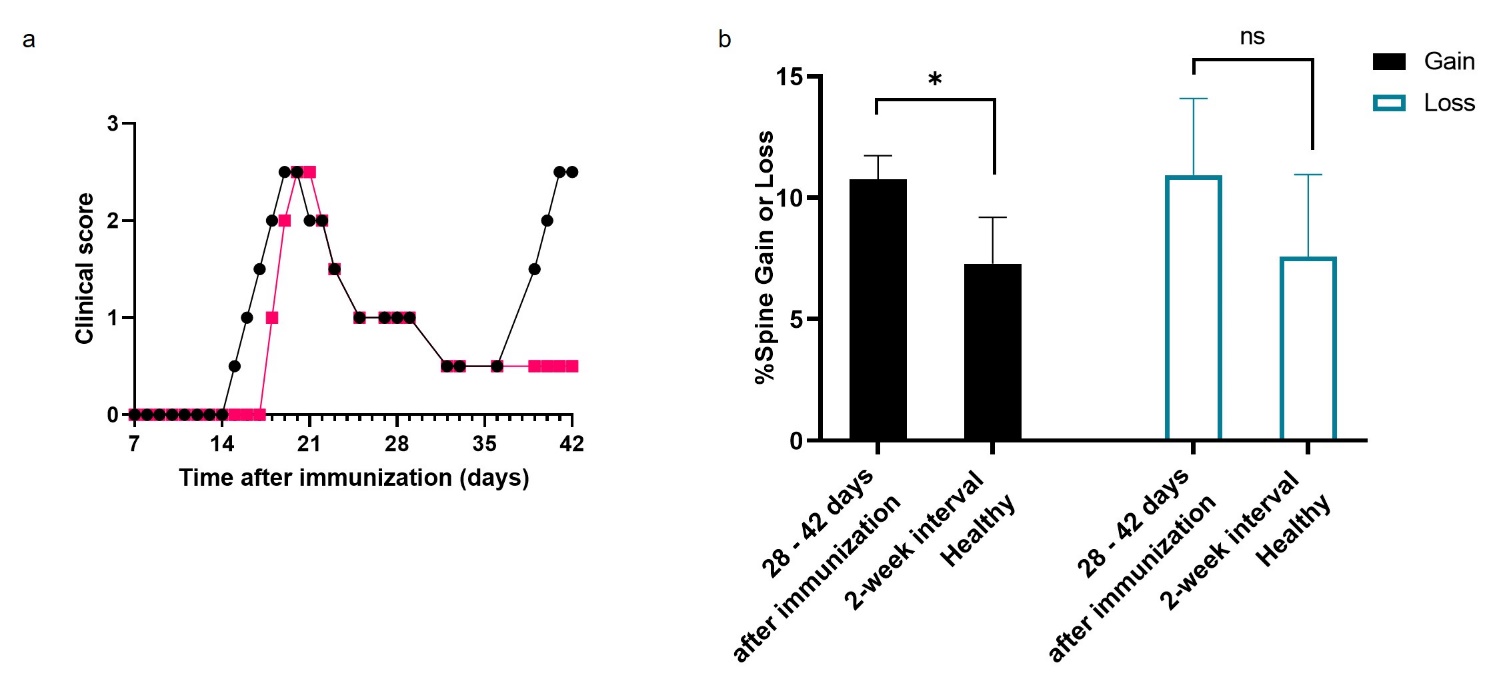

**Supp Fig. 2**

Percent dendritic spines lost or gained from 28 to 42 days after induction of EAE. **a** EAE clinical scores for 2 mice imaged after induction of EAE until 6 weeks after immunization. One of the mice suffered a relapse. Weekly imaging sessions were obtained until 28 days after immunization, and a final imaging session was obtained 2 weeks later. **b** Percent dendritic spines gained or lost. *p = 0.0402 (one-tailed t-test). Mean ± SD. (n = 2 cells from 2 different EAE mice).

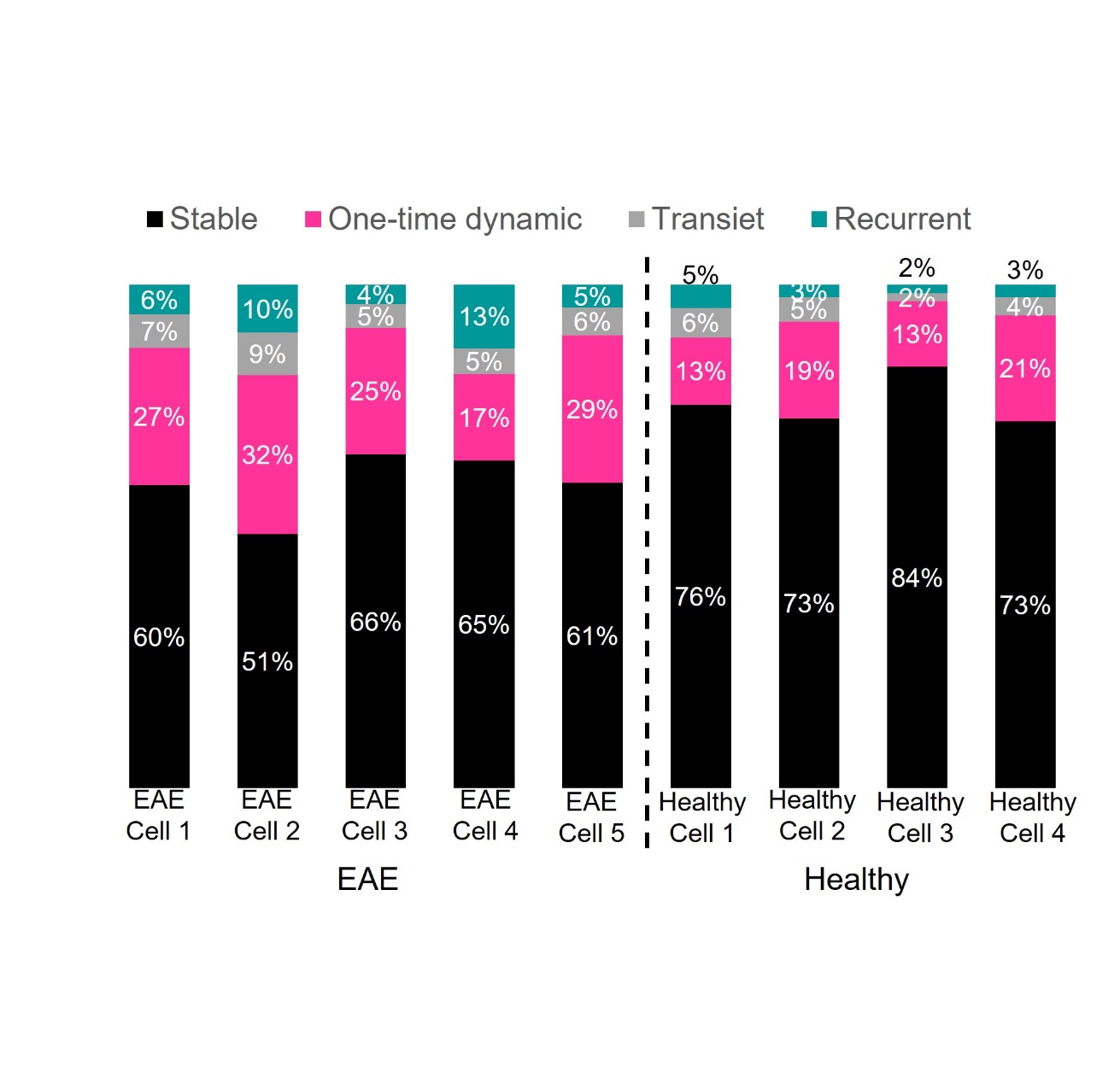

**Supp Fig. 3**

Dynamic behavior of dendritic spines classified as stable, one-time dynamic, transient, and recurrent, as depicted in Fig. 3. Shown are proportions of each category of dendritic spine for the individual imaged cells.

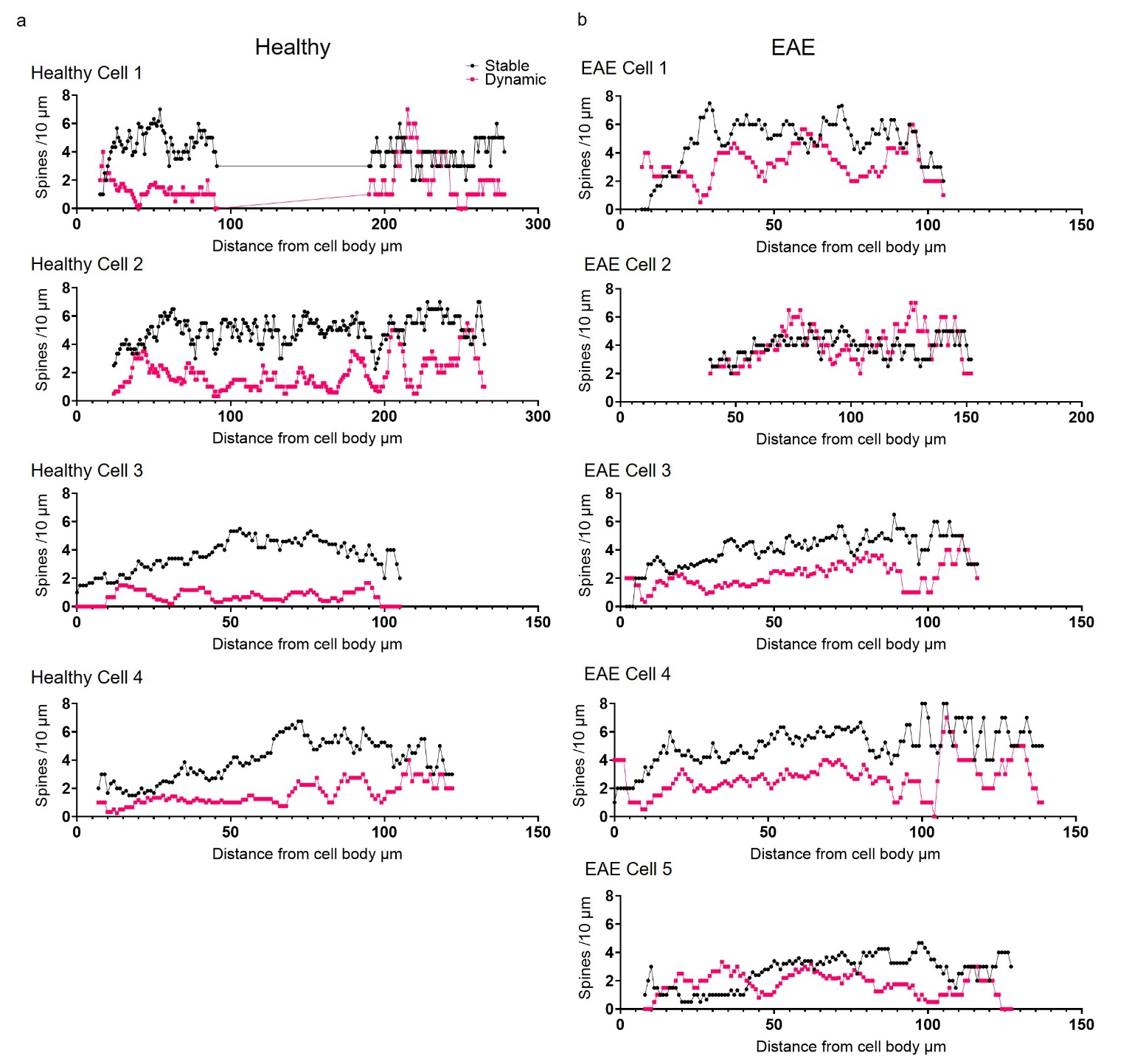

**Supp Fig. 4**

Mean density of dynamic and stable spines for **a** EAE and **b** healthy individual cells by distance from the cell body.
